# Transposase-mediated activity is associated with accessory gene turnover and positive-selection signals in *Streptococcus pyogenes*

**DOI:** 10.64898/2026.11.17.767368

**Authors:** Evan Ho, Omar E. Cornejo

**Affiliations:** Ecology and Evolutionary Biology Department, University of California Santa Cruz, CA, 95064

**Author notes:** Corresponding author: Omar E. Cornejo.

**Keywords:** bacterial pangenome, transposase activity, adaptive evolution, accessory genome

## Abstract

*Streptococcus pyogenes* causes diseases ranging from mild pharyngitis to life-threatening invasive infection, yet the genomic determinants of this variation remain poorly understood. We analyzed 1,863 publicly available genomes, whose pangenome comprises 6,568 genes (1,392 core and soft-core, 656 shell, and 4,520 cloud), and grouped isolates into 15 genetic groups with FastBAPS. Ten groups were monophyletic on the global phylogeny, and non-monophyly of the remaining five largely reflected the placement of isolates from group 12, which is dispersed across the tree. A GenomegaMap scan identified 163 of 3,147 genes with ω₀ > 1. These genes were depleted among core genes and enriched among cloud genes, a pattern consistent with positive selection on recently acquired genes but also with the retention of slightly deleterious variation in young genes. A putative IS transposase (K07497) was the most consistent functional enrichment signal, which led us to analyze insertion sequence (IS) elements directly. In group 12, complete IS copies within genomes were more often near-identical than in other groups (OR = 1.46), consistent with more recent IS activity, and genes with ω₀ > 1 were 2.9 times more likely than other accessory genes to lie within 5 kb of an IS element (8.8% vs. 3.0%; permutation p = 0.0006) after excluding IS-encoded genes. We did not detect this association in any other group, and most genes under selection were not IS-proximal.

## Introduction

*Streptococcus pyogenes* (*S. pyogenes*) causes a wide variety of human diseases, ranging from mild pharyngitis to severe invasive infections such as sepsis and pneumonia (Cunningham 2000; Kanwal and Vaitla 2023). Clinically, *S. pyogenes* has been classified into more than 80 distinct M protein serotypes, a surface protein playing an important role in its virulence (Cunningham 2000). While several *S. pyogenes* outbreaks have been linked to particular serotypes, the prevalence and virulence of others remains unclear (Kaplan et al. 1989). In particular, we still have a limited understanding of whether there are functional differences, mediated by gene content differences, among isolates that are more commonly found in mild infections compared to those found in severe invasive cases.

The factors determining post-infection complications are poorly understood (Walker et al. 2014). For instance, acute rheumatic fever, acute glomerulonephritis, and pediatric autoimmune neuropsychiatric disorder associated with streptococcal infections (PANDAS) are examples of post-infection complications that can occur weeks after initial infection, yet why only some patients develop these complications remains unknown (Orefici et al. 2016; Cocuzza et al. 2022). Differences in the genetic susceptibility of hosts can explain the differential risk to develop these complications, but we have limited knowledge about how much of the genetic variation among bacteria isolates mediates the differential risk to disease. The complex interaction between host and pathogens that ultimately develop in disease is often mediated by genetic differences between hosts as well as genetic differences between bacterial isolates (Didelot et al. 2016; Parks et al. 2020). Understanding the genomic diversity of *S. pyogenes* is a necessary first step toward investigating how pathogenic genetic variation contributes to these differential clinical outcomes.

Bacterial evolution is highly dynamic; genetic information transfers both vertically and horizontally, as well as across species and genera (Clewell 1981). Interspecies horizontal gene transfer among bacteria may contribute to increased resistance to host immunity (Pletz et al. 2006), virulence (Sumby et al. 2005), and transmissibility (Nasser et al. 2014). In *S. pyogenes*, numerous genes have been identified as horizontally transferred from other *Streptococcus* species—including *S. agalactiae* and *S. equi*—and some of these genes have similarity to sequences identified in *Escherichia coli* (Lefébure et al. 2012). Homologous recombination also plays an important role in shaping the genetic landscape and phylogenetic analysis of the *Streptococcus* genus. In *S. mutans*, for instance, recombination played a greater role in the maintenance of genetic diversity than mutation (Cornejo et al. 2013), and in *S. pyogenes* the inferred genetic relationships among strains dramatically shift when corrected for recombination events (Kachroo et al. 2019). In order to better understand the pathogenic potential of *S. pyogenes* and identify subsets of isolates associated with specific disease outcomes, we need to better understand the evolution of the gene composition across a diverse group of isolates.

Several population genomic and pangenomic studies have characterized genomic diversity in *S. pyogenes*, including estimates of core and accessory gene content (Anisimova et al. 2007; Kachroo et al. 2019; Lee and Andam 2022; Jespersen et al. 2024b). In pangenomic analyses, genes are commonly classified as core genes, which are genes present in all or nearly all isolates, and non-core genes. The core genome helps to establish the species’ identity, while non-core genes, also known as shell and cloud genes depending on their frequency, may reveal evolutionary processes that are in flux. Earlier comparative analyses of 26 *Streptococcus* genomes examined recombination and positive selection in the core genome alongside gene gain and loss across the pangenome (Lefébure and Stanhope 2007), while analyses of accessory content focused on species-specific genes and their role in adaptation through gene transfer across 12 genomes from five species (Marri et al. 2006). It remains less clear how recombination and selection act on core, shell, and cloud genes across the species-wide population structure of *S. pyogenes*. Addressing these questions requires integrating pangenomic analyses with phylogenetic structure and evolutionary inference across a large and diverse set of *S. pyogenes* genomes.

This study takes a pangenomic and phylogenetic approach to analyze the genomic diversity and evolution of *S. pyogenes*. We analyzed 1,892 high quality, publicly available annotated genome assemblies from NCBI for our analysis. We identify core, shell and cloud components of S. pyogenes pangenome and use the core genome to evaluate the degree of population structure in the species, and assess the relative impact of selection driving evolution in the core and shell components of the genome. Our results suggest that selection is associated with genes that are disproportionately concentrated within specific groups of isolates. We further identify a transposase-associated signal in the accessory genome and, in a dedicated analysis of insertion sequences, find that complete IS copies are more often near-identical in a single, phylogenetically interspersed group (group 12), where genes with signatures of positive selection also lie closer to IS elements. We interpret these associations as consistent with, but not proof of, a contribution of IS activity to accessory-gene turnover in this lineage.

## Results

### Pangenome composition and gene distribution

To explore the genomic diversity of *S. pyogenes*, an initial pangenomic analysis was performed. Panaroo identified 6,568 genes in the pangenome, composed of 1,284 core genes (∼19.5%), 108 soft core genes (∼1.64%), 656 shell genes (∼9.99%) and 4,520 cloud genes (∼68.8%) (Fig. 1A). This suggests that roughly 21% of the pangenome is indispensable for the species, and 79% may play a role in differentiating isolates into distinct genetic groups. We generated a pangenome accumulation curve by using a rarefaction analysis, and repeated this with 100 permutations to avoid potential sampling biases (Supplementary Fig. 1A). To test whether the *S. pyogenes* pangenome was open or closed, we fitted Heaps’ Law model as well as an asymptotic model to the data to see which was a better fit (Supplementary Fig. 1B). The Heaps’ Law model had a lower AIC score, indicating a better fit, and suggesting that the S. *pyogenes* pangenome is open and has the potential to continuously grow. We also examined gene distribution across isolates and found that a large number of cloud genes are unique to single isolates or shared by only two or three isolates (see Supplementary Fig. 1C). This pattern further supports an open pangenome model.

**Figure 1.**
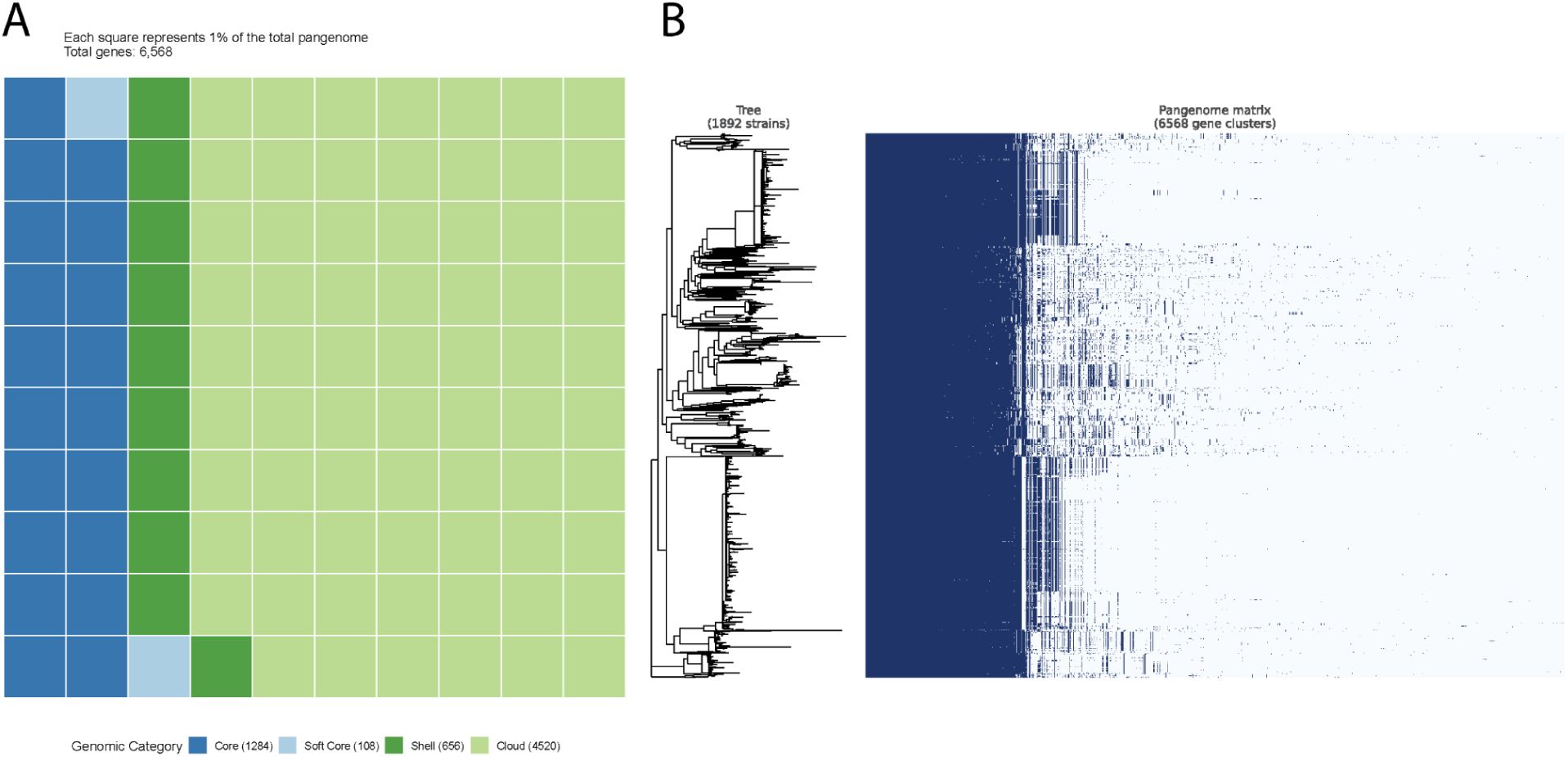
Pangenome composition and its relationship to phylogenetic structure. (A) Distribution of genes across isolates. Genes present in ≥99% of isolates are defined as core, ≥95% and <99% as soft-core, ≥15% and <95% as shell, and <15% as cloud genes. (B) Phylogenetic tree compared to the presence and absence of core and accessory genes. Tree labels are removed for ease of visualization.

In order to examine the phylogenetic distribution of the accessory genome, we then adjoined the RAxML-NG phylogenetic tree with a gene presence-absence heatmap (Fig 1B). Genes are ordered by their frequency across isolates, such that core genes are on the left, followed by shell and cloud genes. This organization facilitates a clearer visualization of gene distribution across the phylogeny and reveals subgroups of isolates with similar gene presence-absence patterns that we further investigate. This visualization provides evidence that some accessory genes are distributed nonrandomly across the phylogeny, potentially reflecting shared evolutionary histories and/or recent horizontal gene transfer events, as well as evidence for within-species genomic diversity.

### Population structure, recombination, and migration among genetic groups

The internal support for some of the observed clades in the phylogeny is relatively small. Because of this we performed a Bayesian hierarchical clustering analysis to infer the number of groups of genetically related isolates in *S. pyogenes*. Our approach provided evidence for 15 groups based on genetic similarity. We used iTOL (interactive Tree of Life) phylogenetic tree visualizer to annotate the phylogenetic tree with the FastBAPS groups (Fig. 2A). Most isolates within a group appear together on the phylogenetic tree, with group 12 being the exception and widely distributed throughout the tree.

**Figure 2.**
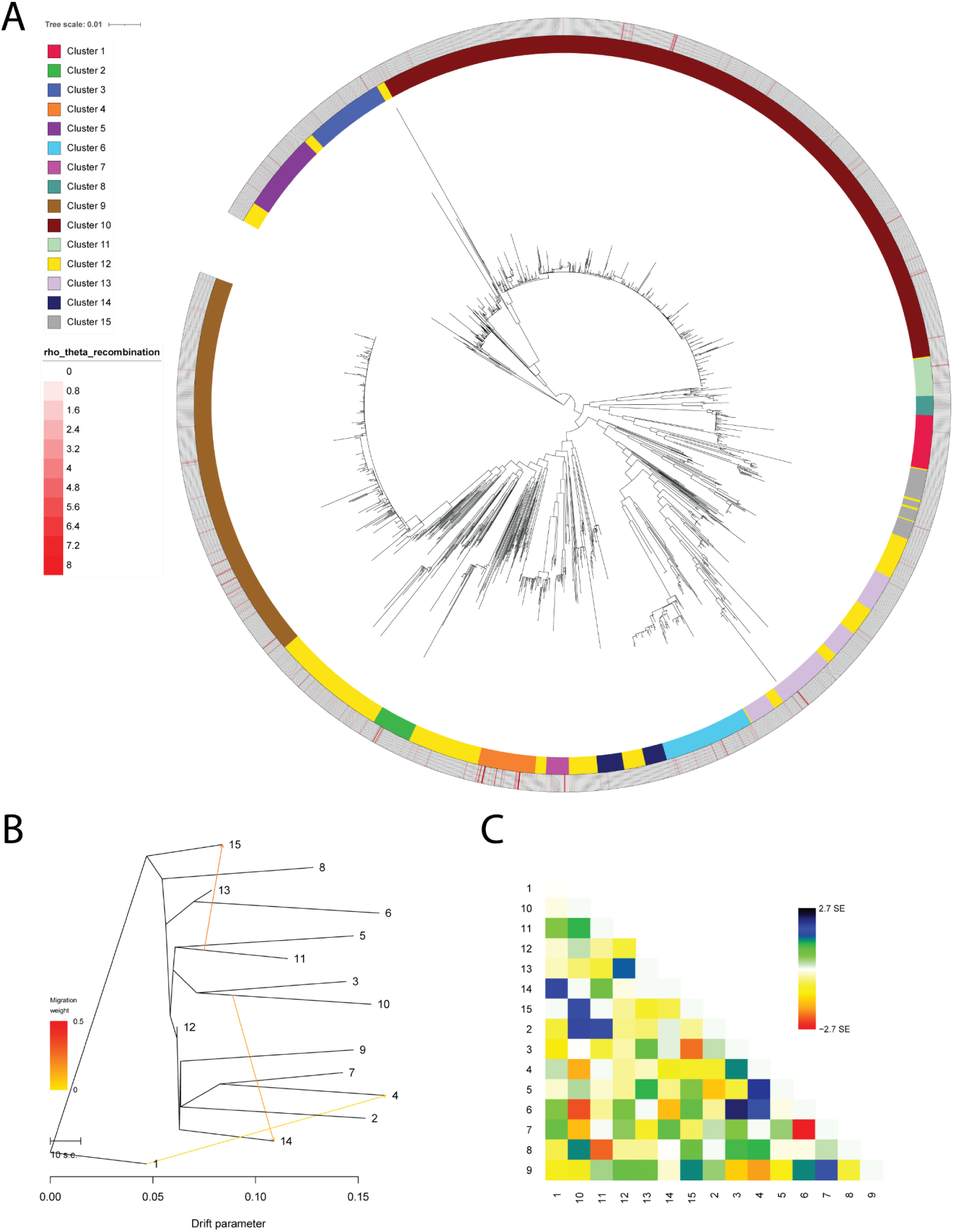
Population structure, recombination, and migration among *S. pyogenes* genetic groups. (A) Phylogenetic structure, grouping, and recombination rates (ρ/θ) of terminal branches, visualized by iTOL. Phylogenetic tree search was performed by RAxML-NG. Bayesian hierarchical clustering of isolates was performed using FastBAPS, which generated 15 groups based on genetically similar isolates (inner circle). Groups are color coded, with the largest group being group 10 (593 isolates) and the smallest group being group 8 (16 isolates). Recombination analysis was performed on each group using Gubbins. (B) Maximum likelihood population tree depicting migration (gene flow) edges among separated genetic groups in *S. pyogenes*. The tree includes three migration events among FastBAPS groups. The x-axis represents the genetic drift parameter, and arrows indicate the inferred migration events, with color intensity reflecting migration weight. (C) Covariance matrix showing the standard error between groups.

To evaluate the robustness of the FastBAPS groupings on the global phylogeny, we assessed monophyly and bootstrap support (200 replicates) for each group. Ten groups (Groups 1-10) were monophyletic on the global phylogeny, each with maximal bootstrap support (FBP and TBE = 100%). Groups 11-15 were not monophyletic: the smallest clade containing all members of each group also contained isolates assigned to other groups (Supplementary Table 1). In groups 11, 14 and 15, these additional isolates were all from group 12 (1, 18 and 5 isolates, respectively). Group 11 was closest to monophyly: its clade contained its 32 members and a single group 12 isolate (GCF_020826955.1). The clade for group 13 contained 144 non-member isolates, comprising all 76 isolates of group 6 (itself monophyletic) and 68 group 12 isolates. Because the tree is unrooted, we repeated these clade assessments under ten alternative rootings (Supplementary Methods; Supplementary Table 2); counts for groups 11, 14 and 15 were unchanged, and the count for group 13 was unchanged unless the tree was rooted within group 6, which lies inside its span. Group 12 isolates were dispersed across the phylogeny: the smallest clade containing all of them included between 1,270 and 1,847 of the 1,863 isolates across the ten rootings, and all 1,863 in the orientation of the RAxML-NG output, where no support value is therefore defined. Because these clades include non-member isolates, support values at their nodes reflect the mixed clades rather than the groups themselves, and we report them in Supplementary Table 1 without interpreting them as support for the groups.

Monophyly is determined from the maximum likelihood tree and therefore does not depend on the number of bootstrap replicates, and groups 1-10 were already at the maximum possible support. Bootstrap convergence, assessed with RAxML-NG’s autoMRE criterion (see Methods), was not reached at 200 replicates (average WRF = 3.11%, 206 of 1,000 permutations at or below the 3% cutoff). Support values for the mixed clades of groups 13-15 changed by no more than 3 percentage points (FBP) and 1.4 (TBE) between 100 and 200 replicates.

We performed recombination analysis using Gubbins within each FastBAPS group separately and generated an annotation file to also display recombination patterns in Fig. 2A. The outer circle displays per-branch ρ/θ values, which is the ratio of recombination events to point mutations inferred by comparing isolates in a phylogenetic context. The combined annotation includes recombination estimates from all 15 group-specific Gubbins analyses mapped onto the global phylogeny. Per-branch ρ/θ and r/m are undefined when a branch has no SNPs outside recombinations, which was true of 1,023 of 3,711 branches (27.6%; Gubbins reports these as 0), and they are unstable when that count is very small. We therefore summarized each group by the proportion of branches with at least one recombination block, which does not depend on this denominator, and by pooled ρ/θ and r/m (total recombination blocks, or SNPs inside recombinations, divided by total SNPs outside recombinations across all branches in the group), with 95% bootstrap confidence intervals. The proportion of branches with at least one recombination block ranged from 22.0% (group 10) to 79.8% (group 12) and was below 50% in 11 of 15 groups. Groups 12 and 7 had the highest proportions (79.8% and 59.5%, respectively) and were the only groups with a non-zero median ρ/θ across all branches (0.058 and 0.071), indicating that recombination was detected on more branches in these two groups. Pooled ρ/θ ranged from 0.0305 (group 14) to 0.2446 (group 2), and pooled r/m from 1.8 (group 12) to 90.8 (group 2) (Supplementary Table 3). We did not compare groups using mean branch-level r/m, because in several groups it is strongly influenced by branches with very few SNPs outside recombinations. Both statistics were right-skewed, with a small number of branches showing very large values (Supplementary Fig. 2).

Among branches with a defined ratio (n = 2,688), groups differed significantly in ρ/θ and r/m (Kruskal-Wallis; ρ/θ H = 252.9, ε² = 0.094; r/m H = 204.4, ε² = 0.076; both p < 0.001), but group membership explained little of the variation among branches (rank-based intraclass correlation 0.106 [95% CI 0.088-0.137] for ρ/θ and 0.085 [0.069-0.114] for r/m), indicating that most variation was among branches within groups. Differences among groups weakened as the minimum number of SNPs outside recombinations increased and were not significant among terminal branches with at least 10 such SNPs (ρ/θ p = 0.060; r/m p = 0.11; Supplementary Table 3). Note that, unlike the iTOL visualization, which shows terminal branches only, these summaries and Supplementary Fig. 2 include all branches.

Because Gubbins was run separately within each FastBAPS group, our ρ/θ and r/m estimates reflect recombination among isolates already assigned to the same group and are not informative about exchange occurring between groups. Consistent with this, patterns of monophyly did not track intra-group recombination rate: groups 2 and 6, which had the highest pooled ρ/θ estimates of the fifteen groups (0.245 and 0.209), were both fully monophyletic with maximal bootstrap support, indicating that substantial recombination can occur within a group without necessarily manifesting as reduced phylogenetic resolution at the group level.

We used TreeMix to infer a maximum likelihood population graph with varying numbers of migration events (indicated by colored arrows). We then defined the model with the number of admixture/gene flow events that maximized the composite likelihood of the inferred configuration, which was three migration events (Fig. 2B). Branch lengths varied among groups, with several groups exhibiting longer terminal branches relative to others. The covariance matrix of the fitted model, which shows the standard error between groups, is reported in Fig. 2C.

### Gene co-occurrence identifies clusters of associated shell genes

Because of the patchy distribution of shell genes along the phylogeny, we performed analyses to identify significant co-occurrence of genes, defined as genes that are found together in clades more often than expected by chance. Coinfinder identified four statistically significant groups of co-occurring shell genes (Fig. 3). In Fig. 3, nodes represent genes and edges represent the strength of the association between them. Most of the genes (N = 637) were assigned to a single large group, while the remaining groups each contained only two or three genes and were omitted from Fig. 3. Although several genes appear to cluster more tightly within the large group, Coinfinder did not classify them as distinct groups. Known shell virulence genes in *S. pyogenes* were highlighted in blue and displayed as slightly larger nodes, and they were all found within the large group, though additional virulence genes exist elsewhere in the pangenome outside the shell gene set shown here. Note that because the Gephi visualization was created using a Fruchterman Reingold method for spatializing the network, genes may be physically closer together yet not statistically significant due to stochasticity.

**Figure 3.**
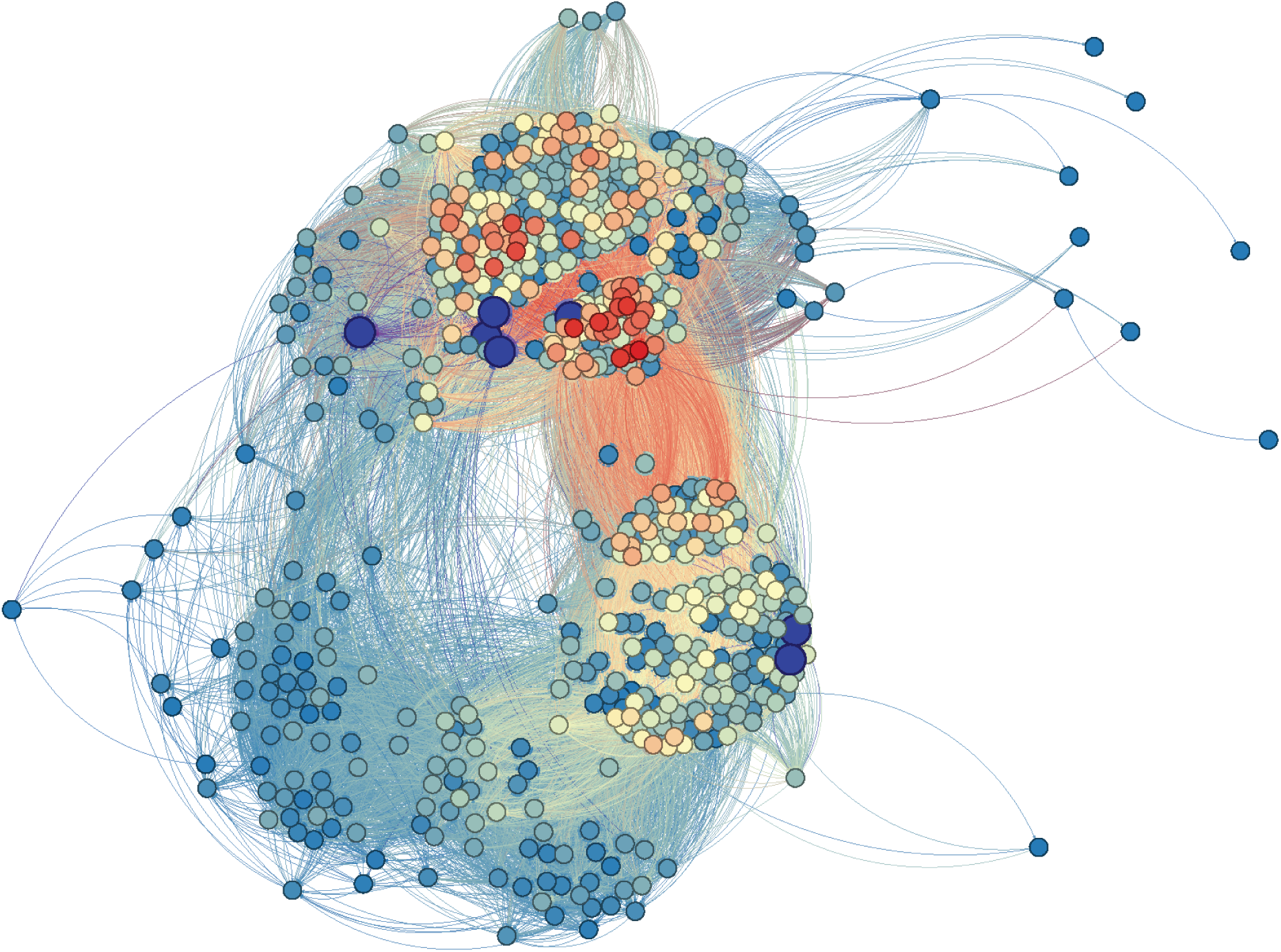
Gephi visualization of significant gene associations. Four groups of significantly associated genes were identified via Coinfinder. The vast majority of genes were assigned to a single large network, shown here; the other three groups were omitted from the visualization because they contained only two or three genes each. Slightly larger blue nodes indicate known shell virulence genes.

### Signatures of positive selection across gene classes

The partition of core and shell genes creates an interesting opportunity to evaluate the impact of natural selection acting on genes that define the species vs. those that potentially define unique functionalities of particular genetic clades within these species. We estimated the evolutionary rate of genes using GenomegaMap, resulting in ω₀, θ, and κ values for each gene (Fig. 4A, Supplementary Fig. 3A–D). Of the 3,439 genes estimated, 3,147 passed Gelman-Rubin convergence diagnostics and were retained for downstream analysis. Both θ and κ distributions had high outliers (θ: 256 genes, 8.1% exceeded the Tukey upper fence, max ≈ 2.8; κ: 121 genes, 3.8%, exceeded the fence, with a small number of genes showing implausibly large estimates, up to ∼4.4 x 10^7^), so multiple visualizations were created to show the distributions with and without outliers.

**Figure 4.**
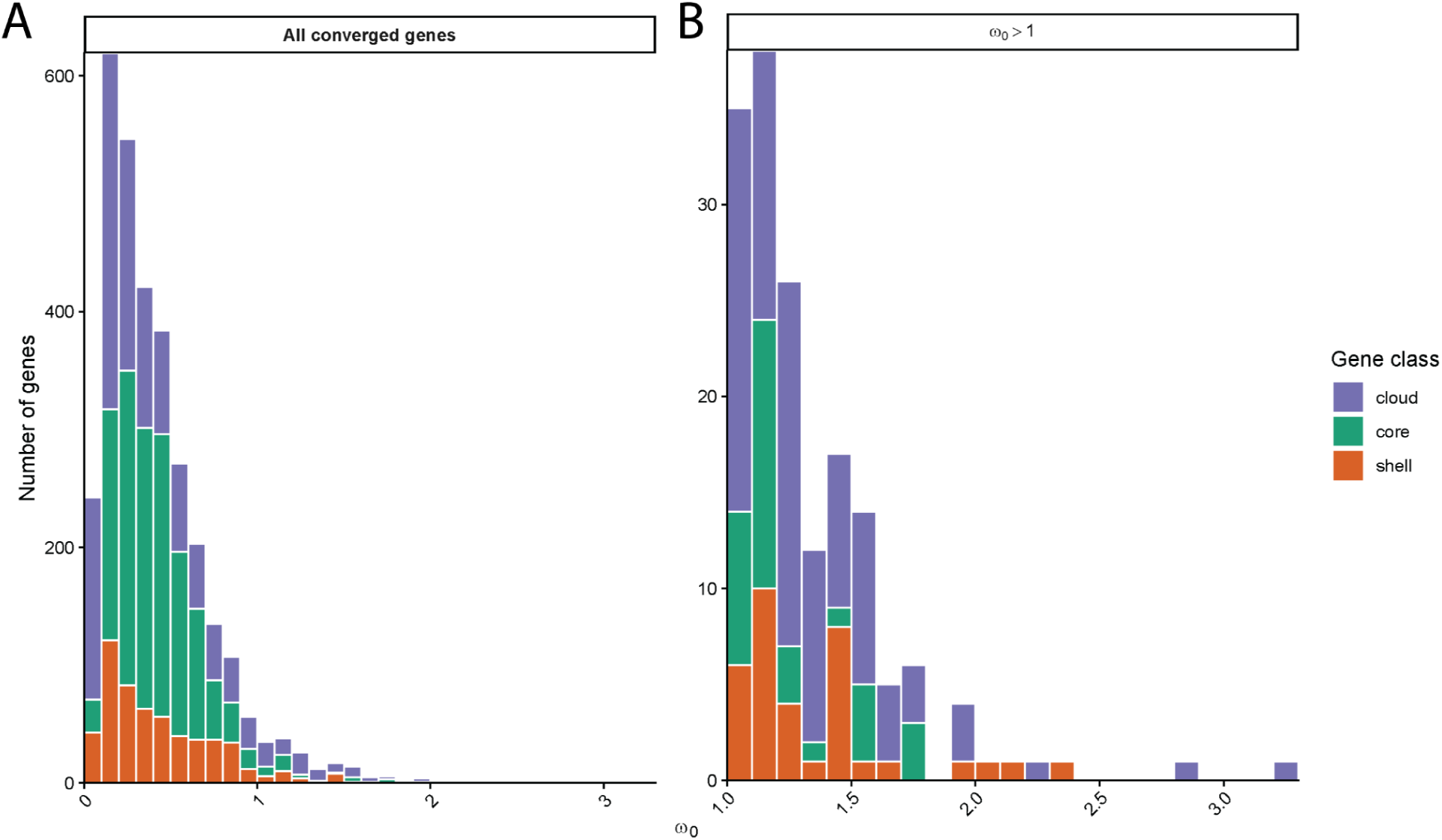
Distribution of ω₀ estimates across genes. (A) All genes. (B) Genes with ω₀ ≥ 1. For each gene, maximum-likelihood parameter estimates from genomegaMap were used to initialize two independent MCMC chains. The first 10,000 iterations were removed as burn-in, and the median ω₀ value was calculated for each chain. The plotted ω₀ values represent the mean of the two chain medians, providing one summary estimate per gene.

Of the 3,147 converged genes, 163 (5.2%) showed evidence of positive selection (ω₀ > 1) (Fig 4B). Of these, 34 (∼21%) were core genes, 35 (∼22%) were shell genes, and 94 (∼58%) were cloud genes. We used a hypergeometric test to assess whether genes under positive selection were over or underrepresented within their respective gene classes, with Benjamini-Hochberg correction applied to account for multiple comparisons across the three gene categories. We found that cloud genes under selection were found at a significantly higher proportion than expected (observed = 94, expected = 62.93, fold-enrichment = 1.49, FDR <0.001), while core genes under selection were found at a significantly lower frequency than expected (observed = 34, expected = 71.02, fold-enrichment = 0.48, FDR < 0.001). Shell genes showed no significant deviation from the null expectation (observed = 35, expected = 29.06, fold-enrichment = 1.20, FDR = 0.127).

### Positively selected genes are concentrated within specific genetic groups

To determine whether positively selected genes were disproportionately associated with particular genetic groups, we tested whether their presence/absence patterns across isolates were more concentrated within specific FastBAPS groups than expected under group-size-proportional random distribution (see Methods). Of the 163 genes with ω₀ > 1, 158 had a valid concentration score (5 were omitted because they were present in all isolates, as concentration could not be assessed for these). Positively selected genes showed significantly higher concentration scores than the remaining converged gene background (median z = 70 vs. 43; Mann-Whitney p = 0.00031; Fig. 5, Supplementary Fig. 4A and 4B), indicating that selection disproportionately acts on genes whose distribution is restricted to particular lineages rather than spread evenly across the population. Of the 158 positively selected genes, 136 (86%) showed individually significant concentration within a specific FastBAPS group (permutation p < 0.05). This signal was highly uneven across groups: FastBAPS groups 12 and 10 alone accounted for 37 and 36 of these significantly concentrated genes, respectively, and together accounting for over half of all significantly concentrated positively selected genes, while the remaining 13 groups each accounted for fewer than 12 genes (Supplementary Fig. 4A).

**Figure 5.**
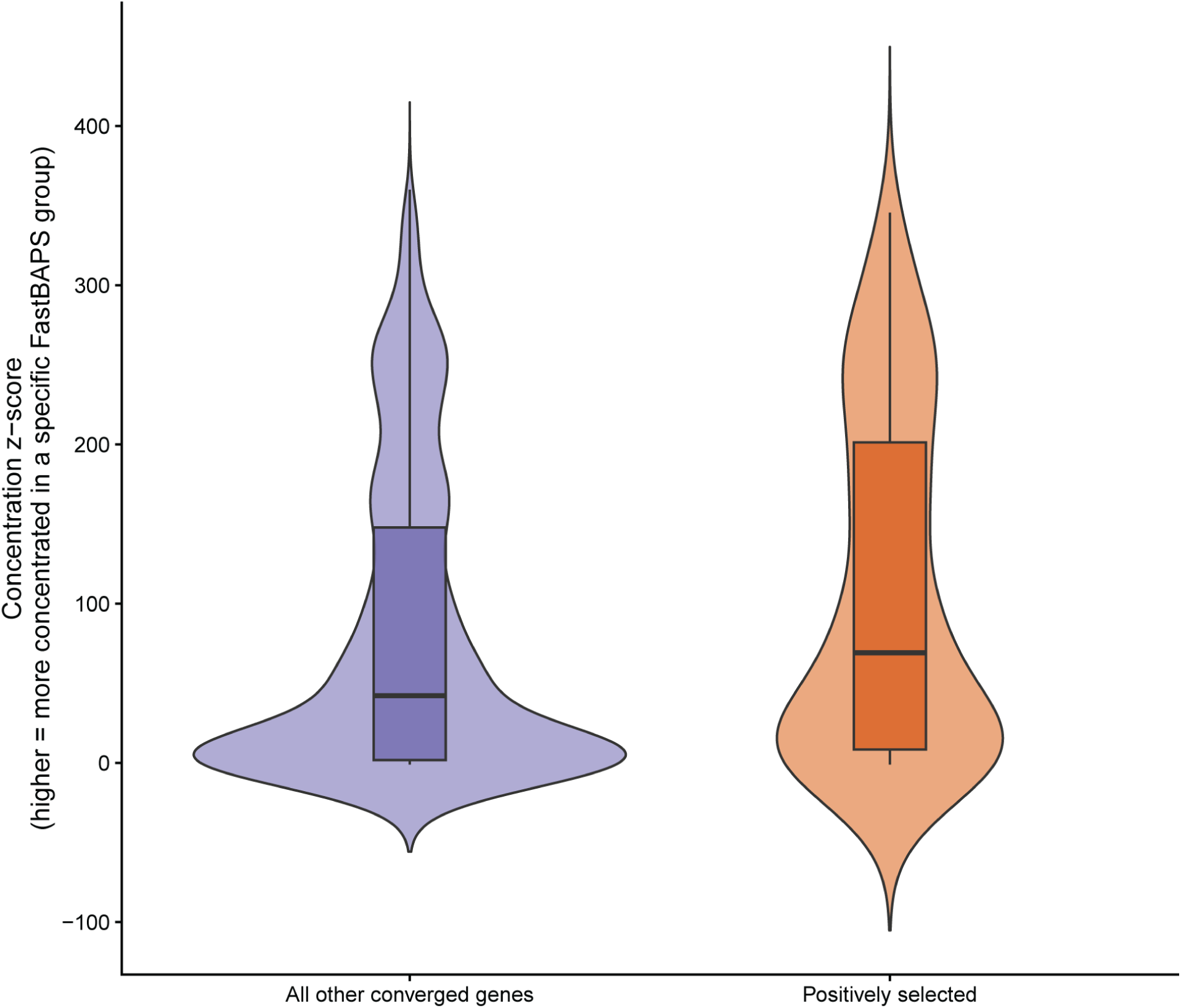
FastBAPS group concentration of positively selected genes relative to the pangenome background. For each converged gene, a permutation-based concentration score (z-score) was calculated reflecting how non-randomly its carriers were distributed across the 15 FastBAPS groups, relative to a null distribution generated from 2,000 random permutations matched for the gene’s carrier count. Positively selected genes (ω₀ > 1; n = 158) show significantly higher concentration scores (median z = 70) than all other converged genes (n = 2,279, median z = 43; Mann-Whitney p = 0.00031), indicating that positively selected genes are disproportionately restricted to specific genetic groups rather than distributed evenly across the population.

However, raw gene counts per group were strongly correlated with FastBAPS group size (isolate count) across all 15 groups (Pearson r = 0.91, p = 2.0 x 10^-6^), and this relationship persisted even after excluding groups 10 and 12 (r = 0.76, p = 0.003; Supplementary Fig. 4C), indicating that larger groups accumulate more raw gene “hits” simply by containing more isolates. When normalized by group size (genes per isolate), this relationship disappeared entirely (r = -0.04, p = 0.89; Supplementary Fig. 4D), and the ranking of groups changed substantially: groups 14 (0.20 genes/isolate) and 11 (0.16) showed the highest normalized concentration, while group 10 – despite its high raw count – fell to below-average density (0.09), consistent with its large size (593 isolates) rather than a distinct biological signal. Group 12 remained among the higher-density groups after normalization (0.13), though no longer the single most concentrated group. Because FastBAPS groups are themselves defined by genetic similarity, this concentration is not, by itself, distinguishable from lineage-restricted genes having been acquired once and subsequently inherited vertically; we address this limitation further in the Discussion.

### Virulence gene prevalence across genetic groups

Next we examined the prevalence of known virulence genes across genetic groups (Fig. 6). Virulence genes were selected based on prior literature establishing their role in *S. pyogenes* pathogenesis (see Supplementary Table 4). Virulence genes were identified across the pangenome in three ways: (1) direct annotation of named Panaroo gene clusters and their presence across isolates; (2) systematic keyword searches of genome annotations for known virulence-associated genes; and (3) BLASTp searches of representative sequences from each alignment against a curated virulence factor database, with hits passing quality control criteria (≥ 70% percent identity, ≥ 70% query coverage and e-value < 1e-10) marked as present. *SfbII_like* represents a collapsed virulence category that contains any genes annotated with fibronectin-binding protein *SfbII*. Some virulence genes are core genes (*arcA*, *tig*, *sagA*, *speB*, *speG*, *slo*, *ska*, *sdaB*, *scpA*, *nga*, and *hylA*), while the distribution of non-core virulence genes is highly heterogeneous; both the distribution of non-core virulence genes across genetic groups and their overall prevalence vary by gene. One known virulence gene, *sib35*, was not detected by any of the three identification approaches used in our pangenome analysis. This absence may reflect limitations in gene clustering by Panaroo and/or failure to meet BLASTp quality control thresholds. We also included a copy number mean and a copy number variance heatmap in the Supplementary Data (Supplementary Fig. 5A and 5B).

**Figure 6.**
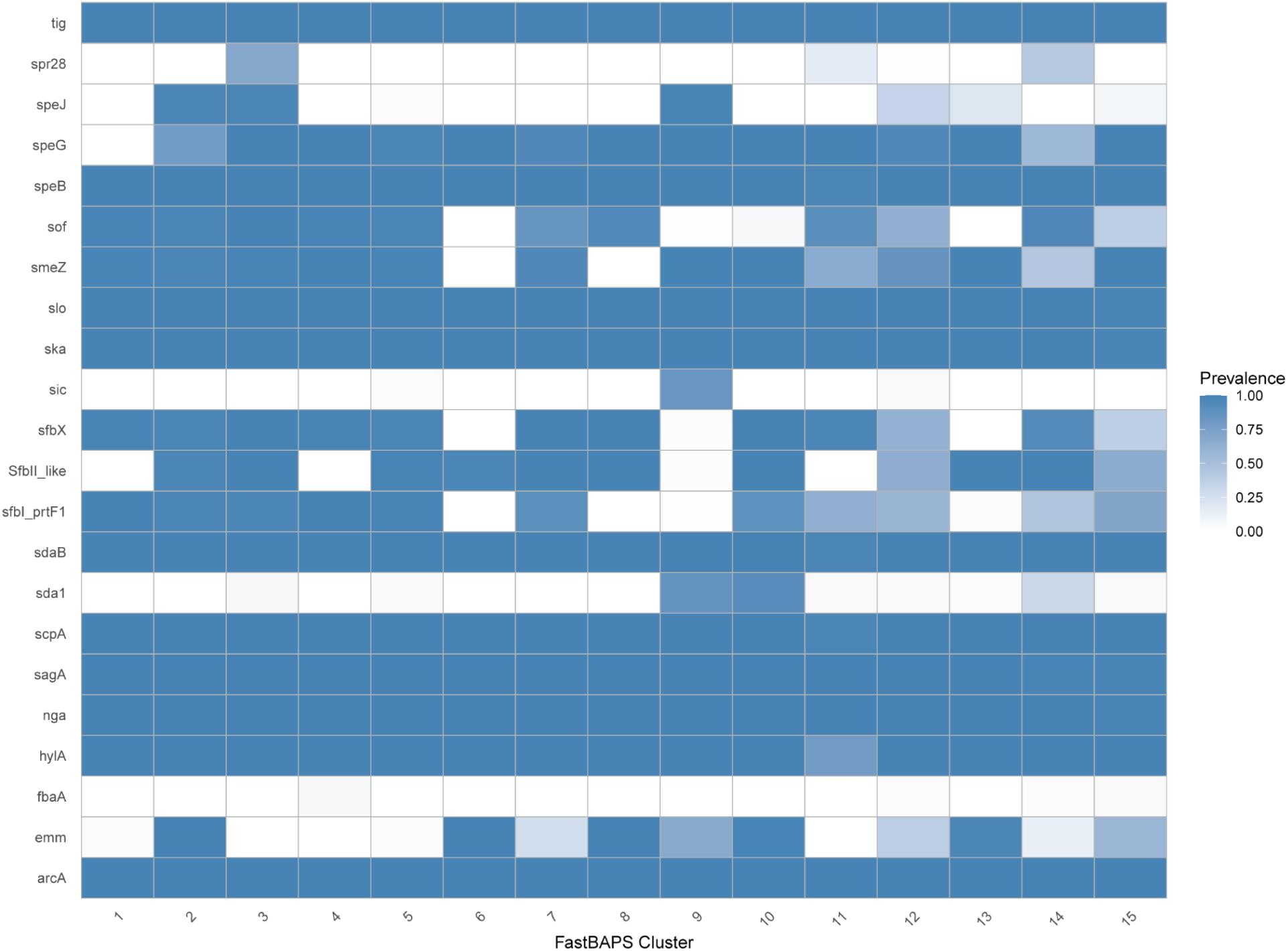
Prevalence of virulence genes across genetic groups. Multiple copies of a gene within a single isolate were counted as a single presence (i.e., gene prevalence was capped at 100%). *fbaA* has low but non-zero prevalence, thus its corresponding heatmap row appears visually uniform despite the presence of a small number of positive isolates.

### Functional enrichment of candidate gene sets

Of the 6,568 pangenome genes submitted to eggNOG-mapper, 477 genes (7.3%) received GO annotations and 2,059 genes (31.3%) received KEGG annotations. Restricting to genes that were both functionally annotated and had converged evolutionary rate estimates, we had 390 background genes for GO enrichment and 1,391 background genes for KEGG enrichment analysis. This background set was reused for enrichment analysis across the three gene sets (all shell genes, Coinfinder-associated genes, positively selected genes). For all shell genes, there were 20 genes for GO and 132 genes for KEGG to test for functional enrichment; 18 and 128, respectively, for Coinfinder-associated; and only 2 and 20, respectively, for positively selected genes. These gene sets are not mutually exclusive (Fig. 7A).

**Figure 7.**
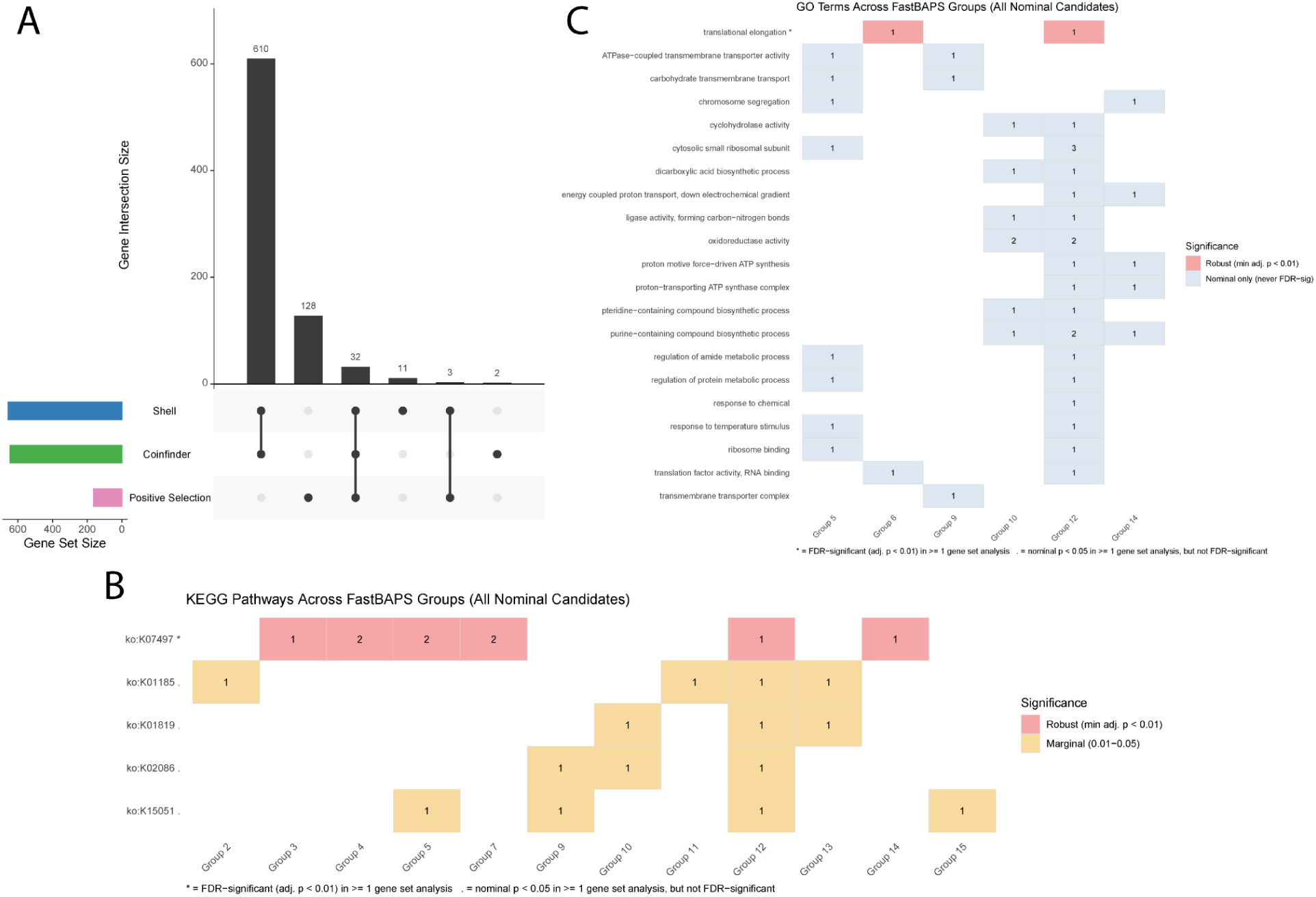
Overlap and functional distribution of candidate gene sets under selection and co-occurrence. (A) Overlap among the three candidate gene sets (shell, Coinfinder-associated, and positively selected genes) prior to restriction to functionally annotated genes. Gene sets are not mutually exclusive; there is substantial overlap between the shell and Coinfinder-associated gene sets, reflecting that Coinfinder co-occurrence analysis was run on the shell genome. Coinfinder-associated and positively selected genes overlap only through genes also present in the shell gene set. (B) KEGG pathway prevalence across FastBAPS groups (all nominal candidates). Tile values indicate the number of genes annotated to each pathway within a group. Color denotes significance tier: robust (★, FDR-significant at adj. p < 0.01 in ≥1 gene set analysis) or marginal (†, nominal p < 0.05 in ≥1 gene set analysis but not FDR-significant). (C) GO term prevalence across FastBAPS groups (all nominal candidates). Tile values indicate the number of genes annotated to each term within a group. Color denotes significance tier: robust (★, FDR-significant at adj. p < 0.01 in ≥1 gene set analysis) or nominal (never FDR-significant in any analysis).

Across all three gene sets, the strongest and most consistent enrichment signal was for KEGG ortholog K07497, annotated as a putative transposase. This term remained significant after FDR correction in every gene set in which it was tested: shell genes (8/132 genes; expected 0.85; adjusted p = 8.06 x 10^-7^), Coinfinder-associated genes (8/128 genes; expected 0.83; adjusted p = 6.27 x 10^-7^), and positively selected genes (4/20 genes; adjusted p = 1.13 x 10^-5^). It is important to note, however, that this is not three independent confirmations of the same result. The genes across shell and Coinfinder-associated test sets are identical, reflecting the substantial overlap between these two gene sets rather than independent evidence. The positive selection gene set contributes a partially overlapping set of 4 genes, 3 of which are shared with the shell/Coinfinder-associated set. Therefore, it is more appropriate to interpret this as a single recurring signal rather than three separate lines of evidence.

Beyond the K07497 result, enrichment was otherwise limited. For shell genes, one additional GO term remained significant after FDR correction: translational elongation (GO:0006414, Biological Process), with 4 of 20 genes annotated to this term against an expectation of 0.37 (adjusted p = 0.007). While no other GO terms passed FDR correction, 17 GO terms have p < 0.05 across the three ontologies (11 Biological Process, 4 Molecular Function, 2 Cellular Component).

For Coinfinder-associated genes, four additional KEGG orthologs have p < 0.05 alongside K07497: endA (DNA-entry nuclease, K15051), lacA_B (galactose-6-phosphate isomerase, K01819), dnaD (DNA replication protein, K02086), and E3.2.1.17/lysozyme (K01185). All four have an identical adjusted p-value of 0.049 in the Coinfinder analysis, just passing FDR correction. That said, the same four orthologs driven by the same genes were also tested in the shell gene analysis (which differs only in test set size: 132 vs 128 genes), where their adjusted p-value is 0.054 and thus do not survive FDR correction. Since this flip in significance is attributable to a four gene difference in the background set, we did not consider these four orthologs robustly significant and report them here as marginal rather than confirmed findings. No GO terms survived FDR correction for the Coinfinder-associated gene set, out of 17 nominal candidates (9 Biological Process, 5 Molecular Function, 3 Cellular Component).

For positively selected genes, the GO test set was too small (2 genes) to detect meaningful enrichment. The KEGG test set (20 genes) yielded only three orthologs with sufficient background representation to test, of which K07497 was the sole candidate.

To determine whether enriched functions were lineage-specific or broadly distributed, we examined their occurrence across FastBAPS groups (Fig. 7B and 7C). Genes carrying the enriched GO terms were heterogeneously distributed, such that only 6 of the 15 FastBAPS groups had enriched GO terms. The FDR-significant shell gene term, translational elongation (GO:0006414), was represented by one gene each in groups 6 and 12. Genes carrying the five candidate KEGG orthologs (K07497 and the four marginal orthologs) spanned 12 of the 15 groups combined. K07497 alone was represented in 6 FastBAPS groups (3, 4, 5, 7, 12 and 14). The three genes detected across all three gene set analyses fell in groups 7, 3 and 4, respectively, while the gene unique to the positive selection analysis was in group 14, a group not shared with any other K07497-carrying gene. The four marginal KEGG orthologs (endA/K15051, lacA_B/K01819, dnaD/K02086, and lysozyme/K01185) collectively spanned 8 groups, 6 of which were not already represented by K07497.

### Transposase activity and its role in gene acquisition

We identified 29,545 putative copies of insertion sequences (IS), out of which 8,414 (28%) were complete. The rest of our analyses were performed only on the complete ISs. We identified an excess of more closely related (higher similarity) ISs in the focal group 12 (Fig. 8A and 8B), which shows more “mobility” in terms of phylogenetic placement (Fig. 8C). The increased odds of observing recent transposition activity in focal group 12 is 1.46 (r2 adjusted p-value = 1.95 × 10⁻⁵). More importantly, these results cannot be explained by quality of assembly alone (see Supplementary Table 5, and Fig. 8C). In group 12, genes under positive selection were more often IS-proximal than other accessory genes (8.8% vs. 3.0% of gene copies; gene-level permutation p = 0.0006). Because neighbouring genes are not independent, we collapsed genes into co-located loci and repeated the test with the locus as the unit. The association persisted (loci containing a selected gene: 9.1% vs. 3.6% of copies within 5 kb of an IS at a 2 kb linkage gap, p = 0.0002; 8.1% vs. 3.8% at 5 kb, p = 0.0016) and was not driven by any single locus (worst-case p = 0.004 after dropping each locus in turn). The association was nonetheless present only in a minority of carrier genomes for most genes: 33 of 86 selected genes had an IS within 5 kb in at least one carrier, but only 5 did in most carriers.

**Figure 8.**
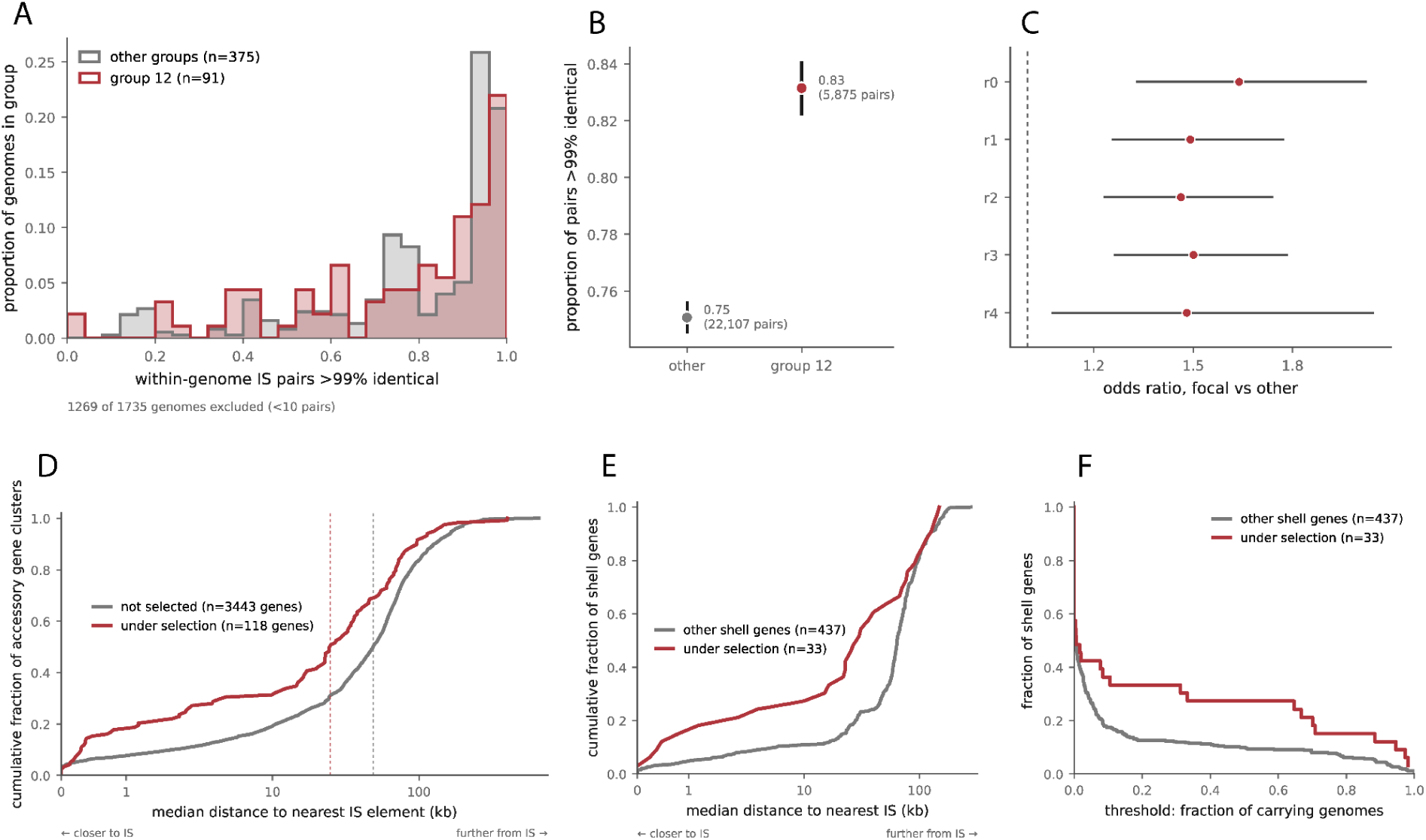
Transposase activity and its association with gene acquisition under selection. (A) Proportion of within-genome ISs >99% identical across all genome comparisons within the focal group (red) and any other group (grey). The proportions are relative frequencies. (B) Point-range plot of the proportion of pairs of IS that are >99% identical in all groups except focal group (grey) and focal group 12 (red). (C) Odds ratios and 95% CI of focal vs. others under different quasibinomial models: r0 presents the unadjusted fit (genomes with more IS copies have more pairs and a higher chance that some pair is near-identical, and group 12 genomes carry slightly more pairs with a median of 5 versus 4); r1 corrects for the total number of pairs and is used as the main model in this comparison, supporting more recent transposase activity in group 12 compared to other groups; r2 is the same model as r1 but adds a correction for assembly fragmentation and is consistent with the main model r1; r3 only corrects for assembly level (no correction for number of pairs) and is used only as a control to understand the impact of assembly level on the estimates; r4 is a more limited model (similar to r1) that constrains the comparisons to only genomes with more than 10 pairs and is used as a control. (D) Cumulative fraction (y-axis) of all accessory genes (shell and cloud) that are under selection (red) and not under selection (grey) as the median distance to the closest IS elements increases (x-axis). (E) Cumulative fraction of only shell genes (y-axis) that are under selection (red) and not under selection (grey) as the median distance to the closest IS elements increases (x-axis). (F) Fraction of shell genes associated with IS-sequences for neutral (grey) and under-selection (red) genes as a function of the fraction of genomes that carry them (the denominator only considers genomes that carry the shell genes). The x-axis is the fraction of genomes carrying IS-associated genes; the y-axis is the fraction of genes that meet a threshold of being within 5 kb of IS elements.

In addition to the increased transposase activity in the more mobile group 12, we found evidence of increased impact of transposase activity in the acquisition of genes under selection in the accessory genome (Fig. 8D and 8E). This effect is not inflated by the larger number of cloud genes, as the cumulative fraction of genes in close vicinity of IS elements is larger in genes under selection with (Fig. 8D) and without (Fig. 8E) cloud genes included. Additionally, if we impose a 5 kb limit on the distance to the closest IS from genes under selection or any other genes within that vicinity of transposase activity, we find that the fraction of shell genes under selection is larger than the shell genes not under selection as we increase the fraction of carrying genomes (Fig. 8F). It is important to note that the denominator in Fig. 8F is limited to the total number of genomes carrying each shell gene assessed. The curve shows the average behavior of the genes under selection (red) and genes that are not under selection (grey). The association was robust to how loci were defined. Across twelve linkage settings (gap of 2 or 5 kb; minimum co-localization in 50% or 80% of carrying genomes; minimum of 3, 10 or 20genomes), the difference in the fraction of copies near an IS between loci containing a selected gene and other loci ranged from 4.3 to 6.9 percentage points, with permutation p-values between 0.0004 and 0.0088 in eleven settings. The exception (p = 0.064) occurred only at the most permissive setting, which merges genes into loci of up to 689 genes (Supplementary Table 6).

## Discussion

*S. pyogenes* genomes are notoriously difficult to characterize due to their highly dynamic nature. Bacterial pangenomes are often open, meaning that with each additional isolate examined, new genes are added to the pangenome. By definition, most of the core genes are captured within just the first few genomes, while the shell and cloud genes are added more gradually as more genomes are added to the analysis. A common question in the analyses of bacterial pangenomes is if they are open, meaning that each additional isolate adds new genes to the pangenome, or if they are closed and have a finite measurable size. One study based on publicly available RefSeq genomes reported that core genes represent ∼5.6% of orthologous genes in *S. pyogenes* (Lee and Andam 2022), whereas another pangenomic analysis of 249 isolates identified ∼24% of genes as core (Jespersen et al. 2024b). These substantial differences may be the result of methodological sensitivity and/or underlying biological heterogeneity within the species, underscoring the complexity of gene presence-absence patterns. Another study focusing on a specific invasive M protein strain (*emm28*) has demonstrated that genetic variation within defined groups can have important functional consequences, including altered gene expression and virulence (Kachroo et al. 2019). Together, these studies highlight the functional relevance of genomic variation, yet leave unresolved how the pangenome is structured across the full *S. pyogenes* population and across different genetic groups.

Our pangenomic analysis of over 1,800 isolates and nearly 6,600 genes revealed several key insights into *S. pyogenes* evolution and diversification. We established a phylogenetic tree of isolates and grouped them by genetic similarity using FastBAPS, which allowed us to analyze the pangenome for recombination and the presence-absence of genes by group. In addition, population relationships among these groups were examined using TreeMix, which identified patterns of genetic relatedness and potential migration events. We also performed extensive selection analyses on shell and cloud genes, extending earlier comparative work on the core genome (Lefébure and Stanhope 2007) and species-specific genes (Marri et al. 2006). We identified enriched functions within and across groups of isolates, suggesting that they may play a role in the diversification.

The pangenomic accumulation curve provides evidence that the pangenome is open, likely a result of their naturally transformable genomes and ability to incorporate DNA from medium (Marks et al. 2014). The phylogenetic tree paired with the FastBAPS groups of isolates reveal that some groups are more genetically diverse than others. Most isolates within group 10, for instance, have short internal branches, as opposed to group 12 which is distributed throughout the phylogenetic tree and has many longer internal branches. The differences in the branch lengths among populations is explained by differences in the magnitude of genetic drift experienced by different populations, as a result of age of the populations, their effective population size, or a combination of both. Additionally, this may indicate that some groups have been the result of more frequent horizontal gene transfer events. To explore this further, our Coinfinder analysis identified 644 shell genes that were significantly co-occurring (of which 637 genes were significantly associating with each other), suggesting that they may be involved in complementary or the same pathways. It is important to note, however, our analysis had 656 shell genes, hence most of the shell genes were identified as co-occurring by Coinfinder analysis.

Although most shell genes were assigned to a single large group in the co-occurrence of genes analysis, there is visually apparent subgrouping within this group (Fig. 3), suggesting that subsets of genes may exhibit stronger associations with one another than with the broader network. Shell genes are not independently distributed across the phylogeny, but instead form a highly interconnected network of genes that tend to be present (or absent) together. This pattern may suggest that groups of genes are inherited together, perhaps via horizontal gene transfer events or that they are closely linked, rather than individually acquired. In contrast, the presence of several small, isolated co-occurring gene groups may represent more discrete functional units that are needed together for them to function, but remain largely independent from the broader shell gene network. Notably, all shell virulence genes were found co-occurring together and within the large group of shell genes, as opposed to forming independent clusters. This organization may reflect functional complementarity or shared evolutionary histories among virulence shell genes. Together, these results suggest that although shell genes exhibit a patchy distribution, they exhibit structured, non-random patterns of co-occurrence in the pangenome.

We found that the proportion of genes under positive selection differed significantly across gene categories, such that core genes were significantly depleted while cloud genes were significantly enriched, and there was not a significant disproportionate presence nor absence of shell genes under positive selection. This suggests that positive selection may be promoting adaptation via selection on newly acquired genes, while the core genome is under strong purifying selection, which is consistent with previous analyses in bacterial systems (Hughes 2005; Hughes et al. 2008).

While we only considered genes that had at least 5 sequences in the alignments, 1,030 genes still did not successfully go though GenomegaMap’s Bayesian inference for parameter estimation, which is about 15.7% of all genes identified in our pangenome. We took the conservative approach to filter out any genes that had at least one negative parameter estimation, though it may be beneficial to investigate this subset of genes using other methods for selection analysis. We also identified a small subset of genes (∼121 genes) that had highly irregular κ values, but decided not to filter those genes out to maintain consistency with only filtering out genes when they had any negative parameter estimates.

We found that positively selected genes were significantly more concentrated within specific FastBAPS groups than expected from the pangenome background (Mann-Whitney p = 0.00031). This concentration was initially most pronounced in groups 12 and 10, but the raw group-level counts were confounded by group size; after normalizing for this (see Results), group 10 no longer stood out as exceptional, while group 12 remained among the more concentrated groups. Because FastBAPS groups reflect genetic similarity and gene content is inherited vertically along lineages, this concentration pattern is consistent both with clade-specific adaptive selection and with the alternative explanation that genes acquired once via horizontal transfer and subsequently retained within a lineage would produce the same signature independent of selection. Distinguishing between these possibilities would require reconstructing the phylogenetic timing of gene gain and loss, which was beyond the scope of this study. We therefore interpret this result as evidence that positive selection andlineage-restricted gene content are associated, rather than direct evidence of group-specific adaptive differentiation.

Our functional enrichment analysis revealed putative transposase (K07497) as the most significantly enriched function, and this signal converges with several other lines of evidence to suggest a broader role for horizontal gene transfer in shaping lineage-specific adaptation in this dataset. A recent pangenomic study found that 118/249 (∼47%) *S. pyogenes* genomes contained at least one chromosomal inversion, indicating that insertion sequence-mediated mobility is a recurrent feature of genome plasticity in this species (Jespersen et al. 2024a). Consistent with this, K07497 was recovered across shell, Coinfinder-associated, and positively selected gene sets, and K07497-carrying genes were distributed across 6/15 FastBAPS groups (3, 4, 5, 7, 12, and 14), suggesting that transposase-associated sequences are recurrently and independently active across multiple lineages rather than reflecting a single clonal expansion event (Siguier et al. 2014).

Group 12 was also notable for its recombination pattern. Recombination blocks were detected on 79.8% of its branches, the highest proportion of any group and far above groups of comparable size (groups 9 and 10: 24.3% and 22.0%). This suggests that recombination is detected across a larger share of group 12’s branches than in any other groups. However, the magnitude of recombination relative to mutation was low: pooled ρ/θ was 0.046 (95% CI 0.043-0.049) and pooled r/m was 1.8 (1.6-2.0), the lowest of any group. Group 12 branches also carry many more SNPs than those of other groups (mean 1,868 per branch versus 135-693), and across groups the proportion of branches with detected recombination increased with SNPs per branch (Spearman ρ = 0.63, p = 0.012), so the breadth of detected recombination in group 12 may partly reflect its branch lengths. Group 12 is also widely distributed across the phylogenetic tree, unlike group 10, whose terminal tips form a single monophyletic cluster. By contrast, group 7 is smaller (37 branches), and its terminal tips form a single monophyletic clade, with a higher recombination magnitude (pooled ρ/θ = 0.191, pooled r/m = 27.3) but a smaller proportion of branches with recombination blocks (59.5%) and no corresponding enrichment of positively selected genes, suggesting that recombination breadth alone doesn’t predict elevated positive selection. Group 12 stands out in the breadth of branches with detected recombination, its phylogenetic dispersion, and the concentration of genes with ω₀ > 1 among its carriers, but not in the magnitude of recombination relative to mutation, which was the lowest of any group. We note that our statistics do not distinguish many independent recombination events from repeated inheritance of fewer ancestral events, so we treat this as a suggestive association rather than evidence of a specific mechanism.

The non-monophyly of the other groups is closely tied to group 12. Group 12 isolates account for all of the non-member isolates nested within groups 11, 14 and 15, and for 68 of the 144 within group 13; the remaining 76 are all of group 6, a monophyletic group lying entirely within the span of group 13 (Supplementary Table 1). Departures from monophyly therefore largely involve the placement of group 12 isolates among other groups, rather than independent disruption of each group, although group 13 additionally encloses all of group 6, which we did not investigate further. This is consistent with group 12’s broad phylogenetic distribution, the high proportion of branches with detected recombination, and elevated positive selection described above. Our data, however, cannot distinguish genetic exchange from the assignment of genetically heterogeneous isolates to a single FastBAPS cluster.

In contrast, groups 1-10 were monophyletic even where within-group recombination was high: groups 2 and 6, which had the highest pooled ρ/θ estimates (0.245 and 0.209), each formed a single clade with maximal support. Given that our recombination estimates cannot detect exchange between groups (see Results), we interpret the dissociation between within-group recombination and monophyly as consistent with genetic exchange that continues readily within some lineages while remaining largely contained by them, potentially reflecting ecological or epidemiological factors that limit gene flow between groups without suppressing it within them.

Enrichment results beyond K07497 were comparatively modest. Four KEGG orthologs in the Coinfinder-associated set (endA/K15051, lacA_B/K01819, dnaD/K02086, and lysozyme/K01185) were marginally significant but did not survive FDR correction in the shell gene analysis, with adjusted p-values shifting from 0.049 to 0.054 due to a small difference in background set size, illustrating a general limitation of FDR-based enrichment testing on moderately sized gene sets. These four orthologs spanned eight FastBAPS groups, six without K07497, hinting that they may reflect a distinct signal from the transposase-driven pattern.

Similarly, the single FDR-significant GO term among shell genes, translational elongation (GO:0006414; adjusted p = 0.007), was driven by only one gene each in two groups (6 and 12) and did not replicate in other gene sets, while GO enrichment for the positively selected set was not meaningfully testable given only two annotated genes.

Our analyses on the transposition activity show an increase in recent transposase activity in isolates of group 12, which is the most “mobile” and phylogenetically interspersed group identified in *S. pyogenes*. Our results are consistent with transposase activity, which is functionally enriched in the shell genome of *S. pyogenes*, contributing substantially to the movement of genetic material across isolates, a result that is unexpected given that *S. pyogenes* is naturally transformable and calls for revisiting the impact that mechanisms alternative to transformation alone have on the evolution of the species and group 12 in particular. In addition to showing elevated transposase activity in a subset of isolates, we find that it is associated with the adaptive acquisition of accessory genes, reflected in the high concentration of genes under selection within the accessory genome of group 12 lineages, and the general evolution of *S. pyogenes*. Together, these results are suggestive that adaptive evolution in this system could be driven by the acquisition of gene clusters that enable lineages to perform functions not previously available to them, opening the possibility of exploring new ecological or host niches. Our results are consistent with IS activity contributing to accessory-gene turnover in group 12, but they do not distinguish this from vertical inheritance, phage or conjugative transfer, or technical artifacts near repeats. Testing this will require insertion-site-resolved comparisons on the phylogeny and long-read assemblies. Despite these caveats, the analyses strongly suggest that the impact of transposase activity in the evolution of bacteria is larger than anticipated.

## Materials & Methods

### Included Streptococcus pyogenes genomes

All annotated *S. pyogenes* genome assemblies available by May 2026 (1,892), were downloaded from the National Center for Biotechnology Information (NCBI). Only annotated genomes that were assembled at the scaffold level or better were included, and atypical genomes were excluded. We also excluded genomes that had a checkM completeness score < 95%. A list of all the included genomes in this study is provided in Supplementary Table 7.

### Pangenome inference

We used Panaroo (Tonkin-Hill et al. 2020) to infer the pangenome and generate unaligned gene sequence fasta files of orthologous gene clusters (full parameters in Supplementary Methods). We translated unaligned gene sequences into protein sequences, aligned them using MAFFT, and used this protein alignment as a scaffold to align the corresponding nucleotide sequences. This is an approach that minimizes reading-frame disruption in downstream selection analyses. We then concatenated the protein-guided nucleotide alignments to build a core genome multiple sequence alignment for phylogenetic and population structure analyses. Genes present in ≥99% of isolates were classified as core, ≥95-99% as soft-core, ≥15-95% as shell, and <15% as cloud genes.

### Genetic grouping using FastBAPS and TreeMix

Phylogenetic tree searches sometimes do not capture the full extent of bacterial evolutionary history (Didelot and Maiden 2010). Therefore, we used FastBAPS to perform Bayesian hierarchical grouping of isolates (Tonkin-Hill et al. 2019). Using the concatenated multiple sequence alignment, FastBAPS automatically generated 15 groups based on genetic similarity, which were then used for further downstream analysis. Furthermore, to investigate the contribution of recombination to the definition of genetic groups, we used TreeMix to infer a maximum likelihood population graph with varying numbers of migration events (Pickrell and Pritchard 2012).

### Assessing recombination within each genetic group

We used Gubbins to assess recombination within each group of isolates and to construct a recombination-filtered phylogeny (Croucher et al. 2015). Because Gubbins requires a syntenic core genome alignment, we first established core gene order using Panaroo’s gene adjacency graph together with Corekaburra, a tool that identifies syntenic blocks of core genes (Jespersen et al. 2022) (full ordering procedure in Supplementary Methods).

Gubbins iteratively identifies loci with elevated densities of clustered base substitutions and constructs a phylogeny based on putative point mutations outside those regions. Running Gubbins separately within each group, and thus on shorter branch lengths, reduces false-positive recombination detection, since longer branches accumulate more substitutions overall, making it harder to distinguish recombination from point mutation.

Recombination-filtered phylogenies were generated using RAxML-NG under a GTR+G model.

### Generating and visualizing a phylogenetic tree

We conducted a phylogenetic tree search using RAxML-NG under a GTR+G model (Kozlov et al. 2019), following RAxML-NG’s built-in quality-control check of the input alignment. This step identified 29 sets of identical sequences; retaining one genome per set removed 29 genomes, reducing the total number of isolates from 1,892 to 1,863. The tree was visualized, unrooted, using the iTOL: Interactive Tree of Life website. A custom script extracted ρ/θ values from the recombination analysis to generate an iTOL annotation file, and the tree was also used to visualize the distribution of core, shell, and cloud genes among isolates.

### Assessing phylogenetic support and monophyly of genetic groups

To evaluate the robustness of the FastBAPS groups on the global phylogeny, we generated 200 bootstrap replicate trees using RAxML-NG and computed both Felsenstein bootstrap proportions (FBP) and transfer bootstrap expectation (TBE) support values (full parameters in Supplementary Methods). Sufficiency of the number of replicates was assessed with RAxML-NG’s autoMRE criterion (3% weighted Robinson-Foulds threshold; full parameters in Supplementary Methods). For each FastBAPS group, we tested whether member isolates formed a monophyletic clade on the unrooted maximum likelihood tree. For monophyletic groups, we extracted bootstrap support at the group’s most recent common ancestor (MRCA) node. For non-monophyletic groups, we identified the smallest clade containing all group members, tabulated the non-member isolates within it by group of origin, and did not interpret support at that node as support for the group.

### Analyzing associations and dissociations between genes

We used Coinfinder to identify genes with significantly associated or dissociated distributions across isolates (Whelan et al. 2020), which groups genes using a Bonferroni-corrected binomial exact test comparing observed to expected co-occurrence rates. Genes present in >95% or <15% of isolates were excluded from this analysis. The resulting gene association network was visualized using Gephi (Bastian et al. 2009) with Fruchterman-Reingold layout.

### Evolutionary rate estimation using GenomegaMap

We used GenomegaMap to estimate within-species genome-wide variation in dN/dS ratio directly from allele frequencies, without requiring a phylogeny (Wilson 2020). After filtering short alignments and sequences with premature stop codons (see Supplementary Methods), 4,469 genes were analyzed using maximum likelihood estimation under a constant per-gene dN/dS model, yielding initial ω₀, θ, and κ parameter estimates. Genes with negative parameter estimates (n = 1,030), which are not biologically interpretable and can result from poor alignments or optimization failure, were excluded from further analysis. The remaining 3,439 genes were analyzed using GenomegaMap’s Bayesian MCMC inference, also under a constant dN/dS model. We assessed convergence between chains using a Gelman-Rubin statistic (full parameters in Supplementary Methods), and 292 genes failed to converge (predominantly shorter genes with fewer aligned sequences) and were excluded, leaving 3,147 genes with fully converged evolutionary rate estimates.

### FastBAPS group concentration of positively selected genes

To test whether positively selected genes were non-randomly distributed across the 15 FastBAPS groups, we compared, for each of the 3,147 converged genes, its observed distribution of presence/absence across groups to a null expectation of group-size-proportional random distribution, using a permutation-based chi-squared approach (full procedure in Supplementary Methods). This yielded an empirical p-value and z-score per gene, allowing concentration to be compared across genes independent of prevalence. This analysis was restricted to the 1,863 isolates with a FastBAPS group assignment and excluded 710 invariant core genes, for which group concentration cannot be meaningfully tested (158 of 163 positively selected genes and 2,279 of 2,984 background genes retained). We compared concentration z-scores between the positively selected gene set (ω₀ > 1) and all other converged genes using a Mann-Whitney U test.

### Functional enrichment analysis

We performed functional enrichment analysis on three gene sets: (1) all shell genes, (2) genes with ω₀ >1, indicating potential positive selection, and (3) genes identified by Coinfinder as significantly co-occurring. For each gene, the longest protein sequence was selected as a representative and functionally annotated using eggNOG-mapper (v2.1.13) (Cantalapiedra et al. 2021) with bacterial taxonomic scope, assigning Gene Ontology (GO) and KEGG orthology terms (full annotation parameters in Supplementary Methods). The enrichment background was restricted to genes that were both functionally annotated and had converged evolutionary rate estimates (see above), computed separately for GO and KEGG given differing annotation success between the two.

GO enrichment was performed using topGO with the weight01 algorithm, which accounts for the hierarchical parent-child structure of the GO graph, using Fisher’s exact test, and was tested separately for the Biological Process, Molecular Function, and Cellular Component ontologies.

KEGG enrichment was performed using clusterProfiler’s enricher() function with a custom term-to-gene mapping and a standard hypergeometric test, without a hierarchy-aware algorithm, as KEGG orthologs lack GO’s hierarchical structure. Each gene set was tested against its corresponding background, restricted to genes from that set present in the relevant background. P-values were adjusted for multiple testing within each ontology/analysis using

Benjamini-Hochberg FDR correction. We report terms/pathways meeting a raw p < 0.05 threshold as candidates, and separately indicate which remained significant after FDR correction; only FDR-corrected results are treated as statistically robust. KEGG orthology term descriptions were retrieved through the KEGG REST API (KEGGREST) in August 2026, and terms without an exact match were cross-referenced manually against the KEGG database (kegg.jp).

### Transposase IS analysis

We analyzed the retained 1863 *S. pyogenes* assemblies. Assembly statistics taken from NCBI metadata were used for the analysis. We performed our analysis with the 15 genetic groups identified with FastBAPS and given the interspersed pattern displayed by group 12 (324 isolates), as well as its accessory genome selection signal, this was designated the focal group of the analysis. Group 12 assemblies were broadly comparable to other groups in contig number, N50, and assembly level composition (Supplementary Fig. 6A-C).

IS elements were identified with ISEScan run on each assembly independently. ISEScan reports terminal inverted repeats (TIR) and target site duplications alongside the transposase ORF, allowing intact elements to be distinguished from degenerate copies; we refer to elements with both TIRs recovered as complete (Xie and Tang 2017). Because short-read assemblies collapse identical repeats and break contigs at them, all IS counts should be read as lower bounds, and the degree of under-counting varies with assembly quality. We considered assembly quality metrics as covariates (Supplementary Fig. 6D), and limit analyses to complete ISs to control for the impact of assembly quality that can inflate estimates.

We modeled IS recency load using a negative quasi-binomial regression with log genome length as an offset and genomes weighted by inverse clonal cluster size. Four nested specifications were fitted: unadjusted; adding contig number and N50; adding assembly level; and restricted to complete copies. Per-family effects were fitted separately with Benjamini-Hochberg correction.

In order to assess how relevant is the impact of recent vs ancient transpositions we computed all-against-all nucleotide identity among its own IS copies within families (BLASTN, ≥70% identity, ≥80% query coverage) for each genome, and recorded the proportion of within-genome pairs exceeding 99% identity. Copies that are near-identical within a single genome cannot have been present since that genome’s core genes diverged, so this proportion indexes recent transposition.

The comparison was made both unadjusted (Mann-Whitney) and as a quasibinomial GLM including contig number and assembly level and total number of IS found. The adjustments matter and runs opposite to the IS-count bias: fragmented assemblies collapse near-identical copies, so poorly assembled genomes appear *less* recently active for purely technical reasons. We show that the quality of the assemblies cannot explain the differences in IS across FastBAPS groups: IS content is not consistently associated with assembly fragmentation and group 12 does not show elevated IS content relative to other groups once stratified by assembly level (Supplementary Fig. 6E and 6F). The main reason to control for the number of pairs found is that if a larger number of IS are found, then there is a higher likelihood to find IS pairs with higher similarity, and this doesn’t necessarily inform recency in the transposase mediated activity.

### Association between IS elements and genes under selection

Accessory genes under selection were taken from our selection scan and ran on FastBAPS group 12 which exhibited a more mobile behavior. We computed the distance from every CDS to the nearest IS element in the focal group, and whether IS elements lay within 15 kb on both sides (a composite-transposon-like configuration). We then compared if the IS were significantly closer to genes under selection than to neutral genes in the focal group.

Distances were aggregated to one median per gene cluster before the analysis, because the raw table contains one observation per gene per genome and treating those as independent inflates the sample size several hundredfold. Significance was assessed by permuting the under-selection label within deciles of accessory-genome frequency (10,000 permutations), because rare genes are both more likely to be recent arrivals and more likely to occupy mobile-element-rich regions. Loci were defined as sets of accessory genes within 2 Kb or 5 Kb of each other in at least 50% or 80% of the group-12 genomes carrying both genes (minimum 3, 10 or 20 genomes), and the locus was used as the permutation unit. We report the range of results across these settings.

For shell genes, we additionally report the proportion of carrying genomes in which an IS element lies within 5 kb, compared between genes under selection and other shell genes (Mann-Whitney on per-cluster proportions), and stratified by gene prevalence.

### Software and reproducibility

Analyses were run on a SLURM-managed high-performance computing cluster in conda environments (Python 3.11 and 3.9; R 4.5.3 and 4.4.3, depending on the pipeline step). Versions of all software and packages, along with the environment and settings used for each step, are listed in Supplementary Table 8.

## Data availability

All genome assemblies analyzed in this study are publicly available from NCBI. GenBank (GCA) and RefSeq (GCF) accessions for all 1,892 downloaded assemblies, with the 1,863 retained after duplicate-genome removal indicated, are listed in Supplementary Table 7. Input and output files for all analyses are archived at Zenodo (https://doi.org/10.5281/zenodo.23403568). The code is also available at https://github.com/evanho550/StrepA.

