## Supplementary_Material for "Transposase-mediated activity is associated with accessory gene turnover and positive-selection signals in *Streptococcus pyogenes*"

### *Pangenome inference*

We used Panaroo (Tonkin-Hill et al. 2020) to infer the pangenome and generate unaligned gene sequence fasta files of orthologous gene clusters, with the following parameters: i) genes must be present in at least 5% of genomes (Panaroo also uses neighborhood information of each node to filter genes, not solely based on gene frequency); ii) initial genes that share a common neighbor are compared at a pairwise sequence threshold of 90%, and collapsed together if they fall within that threshold and annotated accordingly; iii) gene fasta sequences were produced without alignment; and iv) sequences that did not conform to the expected Prokka format, such as those with premature stop codons, were also removed.

We first translated all unaligned gene sequences into protein sequences using a custom script (all custom scripts are provided in the repository). We aligned these protein sequences using MAFFT, then used the protein aligned sequences as a reference to align the gene sequences at the nucleotide level using another custom script. This is an approach that creates gene alignments that will minimize the impact of improper alignment (disruptions of reading frame) on selection analyses. Finally, we concatenated the protein sequence-guided aligned gene sequences to create a multiple sequence alignment of core genes for phylogenetic and population structure analyses.

### *Assessing phylogenetic support and monophyly of genetic groups*

We generated 200 bootstrap replicate trees using RAxML-NG under the same GTR+G model and core genome alignment used for the maximum likelihood tree search. Bootstrap support was mapped onto the best-scoring ML tree using raxml-ng --support, computing both FBP and TBE support values (--bs-metric fbp, tbe). Bootstrap convergence was evaluated post hoc with

RAxML-NG's autoMRE criterion (--bsconverge), using the default cutoff of 3% weighted Robinson-Foulds (WRF) distance (--bs-cutoff 0.03). At 200 replicates the average WRF was 3.11% and 206 permutations were at or below 3%; the test did not converge.

For each of the 15 FastBAPS groups, we evaluated whether member isolates formed a monophyletic clade on the ML tree using the ape and treeio packages in R. For monophyletic groups, we identified the MRCA node of all isolates assigned to that group and extracted the corresponding FBP and TBE support value at that node. For non-monophyletic groups, we identified the smallest clade containing all members, tabulated non-member isolates by group of origin, and did not interpret support at that node as support for the group. MRCA support extraction was independently verified for the largest group (group 10) using ete3 to confirm consistency between implementations.

Because the tree is unrooted, clade compositions depend on its orientation. We therefore re-rooted the tree on one isolate from each of Groups 1-10 and recomputed the compositions for Groups 11-15 (Supplementary Table 2). For Groups 11, 14 and 15 the number of non-member isolates was identical under all ten rootings (1, 18 and 5, respectively). For Group 13 it was 144 under nine rootings; rooting on a Group 6 isolate, which places the root within the span of Group 13, gave 1,661. The smallest clade containing all of group 12 isolates varied between 1,270 and 1,847 isolates across the ten rootings, compared with all 1,863 in the orientation of the RAxML-NG output.

### *Core genome gene ordering for recombination analysis*

To use Gubbins to estimate recombination across phylogenetic groups, we first established the syntenic order of core genes. Core genome gene order was established using a combination of Panaroo's gene adjacency graph and Corekaburra, a tool that identifies syntenic blocks of core genes (Jespersen MG et al. 2022). Genes belonging to the same Corekaburra syntenic block were grouped together and treated as a single ordered unit, since their internal order was already established by Corekaburra; core genes not part of any syntenic block were treated as individual units. Together, these blocks and individual genes formed 611 ordering units spanning the pangenome.

To arrange these units into a full genome order, we constructed a second graph in which each unit was a node, connected to other units based on Panaroo's gene adjacency data and weighted by the number of genomes supporting each adjacency. This graph consisted of 23 segments; within each segment, we began at the weakest-supported connection (treated as the natural break point in the circular chromosome) and built the gene order by repeatedly moving to the most well-supported neighboring unit, restarting from any unplaced unit if a dead end was reached. The direction of each syntenic block (forward or reverse) was determined by which end of the block connected to the preceding unit. This process produced a final ordered list of all 1,392 core genes, combining Corekaburra-defined syntenic blocks with individually placed genes, which was used for downstream core genome alignment.

### *GenomegaMap filtering and convergence assessment*

Cloud genes with fewer than 5 sequences in their alignment were removed prior to analysis, excluding 2,099 genes. GenomegaMap also requires removal of sequences with premature stop codons; while these are typically filtered by Panaroo, Panaroo's gene recovery step does not apply this filter, so a custom script was used to remove any remaining sequences with premature stop codons from all gene alignments (see Supplementary Data for the number of sequences removed per gene alignment).

MCMC-based Bayesian parameter estimation was run as two independent chains of 100,000 iterations per gene. Convergence was assessed using a custom script to compute a Gelman-Rubin statistic, which quantifies between-chain variance relative to within-chain variance. The first 10,000 iterations of each chain were discarded as burn-in prior to this calculation.

### *FastBAPS group concentration permutation test*

For each gene, we determined its carrier count (number of isolates carrying that gene) and calculated the expected number of carriers per FastBAPS group under a null hypothesis of group-size-proportional random distribution. We compared this to the observed carrier count per group using a chi-squared statistic, computed as the sum across all 15 groups of the squared difference between observed and expected carrier counts, divided by the expected count. To generate a gene-specific null distribution, we randomly sampled without replacement the same number of isolates as the gene's carrier count from the full isolate pool, 2,000 times, recalculating the chi-squared statistic for each permutation. Each gene's observed statistic was compared to its permuted null distribution to generate an empirical p-value and z-score.

### *Prevalence of virulence genes across FastBAPS groups*

The list of candidate virulence genes analyzed in this study was compiled through a review of the published literature on *S. pyogenes* virulence and pathogenesis (Supplementary Table 4). For the BLASTp-based identification approach, representative sequences were queried against a curated *S. pyogenes* virulence database built in-house.

*Functional annotation and enrichment parameters*

Protein sequences were annotated using eggNOG-mapper (v2.1.13) in DIAMON search mode, using DIAMOND’s default sensitivity setting (sensitive, with adaptive per-query iteration enabled by default). GO enrichment testing used a minimum node size of 5 genes.
